# Direct comparison of contralateral bias and face/scene selectivity in human occipitotemporal cortex

**DOI:** 10.1101/2021.05.06.442603

**Authors:** Edward H Silson, Iris I A Groen, Chris I Baker

**Author notes:** equal contribution (Edward H Silson, Iris I A Groen).

## Abstract

Human visual cortex is organised broadly according to two major principles: retinotopy (the spatial mapping of the retina in cortex) and category-selectivity (preferential responses to specific categories of stimuli). Historically, these principles were considered anatomically separate, with retinotopy restricted to the occipital cortex and category-selectivity emerging in the lateral-occipital and ventral-temporal cortex. However, recent studies show that category-selective regions exhibit systematic retinotopic biases, for example exhibiting stronger activation for stimuli presented in the contra- compared to the ipsilateral visual field. It is unclear, however, whether responses *within* category-selective regions are more strongly driven by retinotopic location or by category preference, and if there are systematic differences *between* category-selective regions in the relative strengths of these preferences. Here, we directly compare contralateral and category preferences by measuring fMRI responses to scene and face stimuli presented in the left or right visual field and computing two bias indices: a contralateral bias (response to the contralateral minus ipsilateral visual field) and a face/scene bias (preferred response to scenes compared to faces, or vice versa). We compare these biases within and between scene- and face-selective regions and across the lateral and ventral surfaces of visual cortex more broadly. We find an interaction between surface and bias: lateral surface regions show a stronger contralateral than face/scene bias, whilst ventral surface regions show the opposite. These effects are robust across and within subjects, and appear to reflect large-scale, smoothly varying gradients. Together, these findings support distinct functional roles for lateral and ventral visual cortex in terms of the relative importance of the spatial location of stimuli during visual information processing.

## Introduction

Visual cortex in each hemisphere initially receives visual inputs from different parts of the visual field, whereby the left visual field is mapped onto the right hemisphere and the right visual field to the left hemisphere (Wandell et al. 2007). This contralateral mapping of visual inputs is the most fundamental organizational feature of bottom-up visual processing in visual cortex, with cross-talk between hemispheres presumably requiring connections across the corpus callosum (Wandell et al. 2007). Within each hemisphere, incoming visual input continues to be processed in a spatially licensed manner: nearby points in the visual field are processed by receptive fields at nearby locations on the cortical sheet. This systematic spatial mapping of visual inputs is known as *retinotopy*, a major organising principle of visual cortex that is commonly used to subdivide cortex into a series of maps (Wandell et al. 2007) that are thought to give rise to a cortical hierarchy consisting of distinct visual areas.

Another key organising principle of visual cortex is *category-selectivity*, which describes the phenomenon that some brain regions respond more strongly to the sight of specific stimulus classes, such as faces, scenes, and objects, compared to others (Kanwisher and Dilks 2013). Category-selective regions were originally identified in cortical locations more anterior to the first retinotopic maps in the hierarchy (V1, V2 and V3). As a result, these two organising principles have historically been thought of as anatomically separate, with retinotopy considered predominant in posterior, early visual cortex (EVC) and category-selectivity considered predominant in the relatively more anterior, lateral-occipital cortex (LOTC) and ventral-occipitotemporal cortex (VOTC), respectively (Op de Beeck et al. 2019). However, subsequent studies revealed that category-selective regions are sensitive to visual field position akin to retinotopic regions (Levy et al. 2001; Hasson et al. 2002). In addition, systematic comparisons of higher-order retinotopic maps and category-selective regions show considerable overlap (Larsson and Heeger 2006; Sayres and Grill-Spector 2008; Arcaro et al. 2009; Silson et al. 2016).

In particular, consistent with the contralateral mapping of visual inputs into the brain, object-scene-, body- and face-selective regions in each hemisphere show a preference (i.e. stronger response) when stimuli are presented in the contralateral visual field (Hemond et al. 2007; MacEvoy and Epstein 2007; Chan et al. 2010, Uyar et al., 2016). While neuronal responses in higher-level visual regions are generally found to be more tolerant to stimulus position than neurons in early visual cortex, the responses are not entirely position invariant (Hong et al. 2016; Apurva Ratan Murty and Arun 2018), and position information can be decoded from fMRI responses in category-selective regions (Schwarzlose et al. 2008; Kravitz et al. 2010; Carlson et al. 2011). Collectively, these findings demonstrate that in addition to a *category preference*, category-selective regions in LOTC and VOTC also contain a *spatial preference* for information from specific parts of the visual field. Indeed, recent population receptive field (pRF) mapping experiments by our group (Silson et al. 2015) and others (Kay et al., 2015; Gomez et al. 2018) demonstrate that category-selective regions throughout LOTC and VOTC exhibit reliable retinotopic biases with a consistent bias for the contralateral visual field. There are also systematic differences in retinotopic preference between LOTC and VOTC, with regions in LOTC exhibiting a lower field bias and regions in VOTC exhibiting an upper field bias (Silson et al. 2015), perhaps reflecting different functional roles (Kravitz et al. 2010).

Despite demonstrating the co-localization of retinotopy and category-selectivity throughout visual cortex, prior work has not directly compared the relative strength of these two factors within regions. That is, although both factors have been shown in, for example, the occipital place area (OPA), it is unclear whether its category bias for scenes (over faces) is greater than its bias for stimuli in the contralateral (over ipsilateral) visual field. Identifying the relative strength of these organizational principles within category-selective regions is an important step towards understanding how the representation of visual space and object identity interact in the brain (Uyar et al. 2016). Here, we investigate the relative strength of retinotopy and category-selectivity directly by presenting face and scene stimuli to either the left or right visual field, thereby making the stimuli exclusively available (initially) to one hemisphere at a time. We chose these particular categories and visual field positions because these provide the strongest possible test of interaction between category and visual field positions: visual cortex is known to contain multiple face and scene preferring regions with divergent and strong preferences between these categories (Julian et al. 2012; Weiner et al. 2018; Margalit et al. 2020), and the difference between contralateral and ipsilateral visual fields provides the strongest retinotopic effects.

This paradigm allows us to compute two bias indices: a contralateral bias (response of stimuli in the contralateral minus ipsilateral visual field) and a face/scene category bias (preferred response to faces compared to scenes, or vice versa). We first compare these biases in independently localized scene- and face-selective regions in both LOTC and VOTC. We then characterize these biases more broadly across the cortex, revealing qualitatively different gradients between LOTC and VOTC, respectively. Together, our results reveal the presence of an interaction between bias (contralateral vs. face/scene) and cortical surface (lateral vs. ventral), resulting in ventral face- and scene-selective regions showing a more pronounced face/scene bias than contralateral bias, whilst lateral regions show the opposite pattern.

## Methods and Materials

### 1. Participants

A total of 18 participants completed the experiment (14 females, mean age = 24.8 years). All participants had normal or corrected to normal vision and gave written informed consent. The National Institutes of Health Institutional Review Board approved the consent and protocol. This work was supported by the Intramural Research program of the National Institutes of Health – National Institute of Mental Health Clinical Study Protocols 93-M-0170 (NCT00001360) and 12-M-0128 (NCT01617408).

### 2. Overview of experimental design

Each participant completed four fMRI sessions: an initial functional localizer session, followed by three independent experimental sessions. In the experimental sessions, participants were presented with four runs of the lateralized scene-face paradigm (see below), and then removed from the scanner to receive theta-burst stimulation to either scene- or face-selective regions of interest, after which scanning was resumed immediately (consecutive TMS-fMRI paradigm). For the purpose of the current study, we only analyzed the pre-TMS runs. Images were repeated across pre-TMS runs in all three experimental sessions, but participants always saw a different set of images in the post-TMS runs. Therefore, our results do not include any potential effects of TMS.

### 3.3. OT scanning parameters

All functional data were acquired on a 3.0T GE Sigma MRI scanner in the Clinical Research Center on the National Institutes of Health campus (Bethesda, MD). Whole-brain volumes were acquired using an eight-channel head coil (28 slices; 3×3×4mm; 10% interslice gap; TR, 2 s, TE, 30ms; matrix size, 64×64, FOV, 192mm). T1-weighted anatomical images were acquired using the magnetization-prepared rapid gradient echo (MPRAGE) sequence (176 slices; 1×1×1mm; TR, 2.53 s, TE, 3.47 ms, TI, 900 ms, flip angle 7°) in the localizer session and in each TMS-fMRI session both before and after TMS.

### 4. Visual stimuli and task

#### 4.1 Functional localizer session

This session consisted of six category localizer runs during which color images from six categories (Scenes, Faces, Bodies, Buildings, Objects and Scrambled Objects, 768 × 768 pixels, 240 exemplars per category). Images were collected from prior experiments run in our lab supplemented with images sourced from the internet and self-taken photos. Scene images were equally divided between indoor, outdoor man-made and outdoor natural scenes (80 images each). Face images were taken from frontal viewpoints and were balanced for gender (120 male, 120 female); moreover, care was taken to introduce variety in race, hairstyle, etc. Bodies consisted of pictures of hands (120 images) and feet (120 images) taken from a variety of viewpoints. Buildings consisted of a large variety of human-built structures (including houses, apartment buildings, arches, barns, mills, towers, skyscrapers, and so on). Objects consisted of both man-made items (120 images, including, amongst other things, household items, vehicles, musical instruments, electronics and clothing) and natural items (120 images, including, amongst other things, fruits/vegetables, nuts, rocks, flowers, logs, leaves, and plants). Faces, bodies, buildings and objects were cropped out and placed on grayscale backgrounds. Scrambled images were created by taking the cropped object images and randomly swapping 48×48 pixel ‘blocks’ across images. Image exemplars were randomly sampled during presentation, but stimulus selection was constrained such that subcategories (e.g. gender for faces, man-made/natural for objects) were equally often presented in each run. Stimuli were presented at fixation in 16 s blocks (20 images per block, 300 ms per image, 500 ms blank). We chose to present the localizer stimuli centrally because this is how category regions are typically mapped. Images were back-projected on a screen mounted onto the head coil with 1024×768 pixel resolution and presented at 10×10° degrees of visual angle). Blocks were separated by 4 s blanks and started and ended with a 16 s baseline period. The total run length was 279 seconds. Each category was presented twice per run, with the order of presentation counterbalanced across participants and runs. Participants performed a one-back task on the images, with 1-3 repeats per block.

### 4.2 Lateralized scene and face sessions

Participants fixated a central cross whilst colour images (5×5° visual angle) of scenes and faces. There were 80 exemplars per category, selected randomly from the larger set of images used in the localizer experiment, whilst ensuring an equal proportion of male/female faces and indoor/outdoor man-made/outdoor natural scenes. Stimuli were presented to either the left or right visual field, centered at 5° offset from the screen center, creating a gap of 2.5° on either side of the fixation point (**Figure 1B**). Images were back-projected on a screen mounted onto the head coil with 1024×768 pixel resolution. Images were presented in 16 s blocks (20 images per block, 300 ms per image, 500 ms blank). Consecutive blocks were separated by 8 s blank periods; in addition, each run started with and ended with a 16 s blank baseline period and included a 16 s baseline period in the middle of the run, resulting in a total run length of 415 s. As each stimulus was presented, one arm of the fixation cross (either horizontal or vertical) increased in length. Participants were required to identify, via button response, the longer arm. Stimulus presentation and fixation cross changes occurred simultaneously. Accuracy and reaction times were recorded.

**Figure 1:**
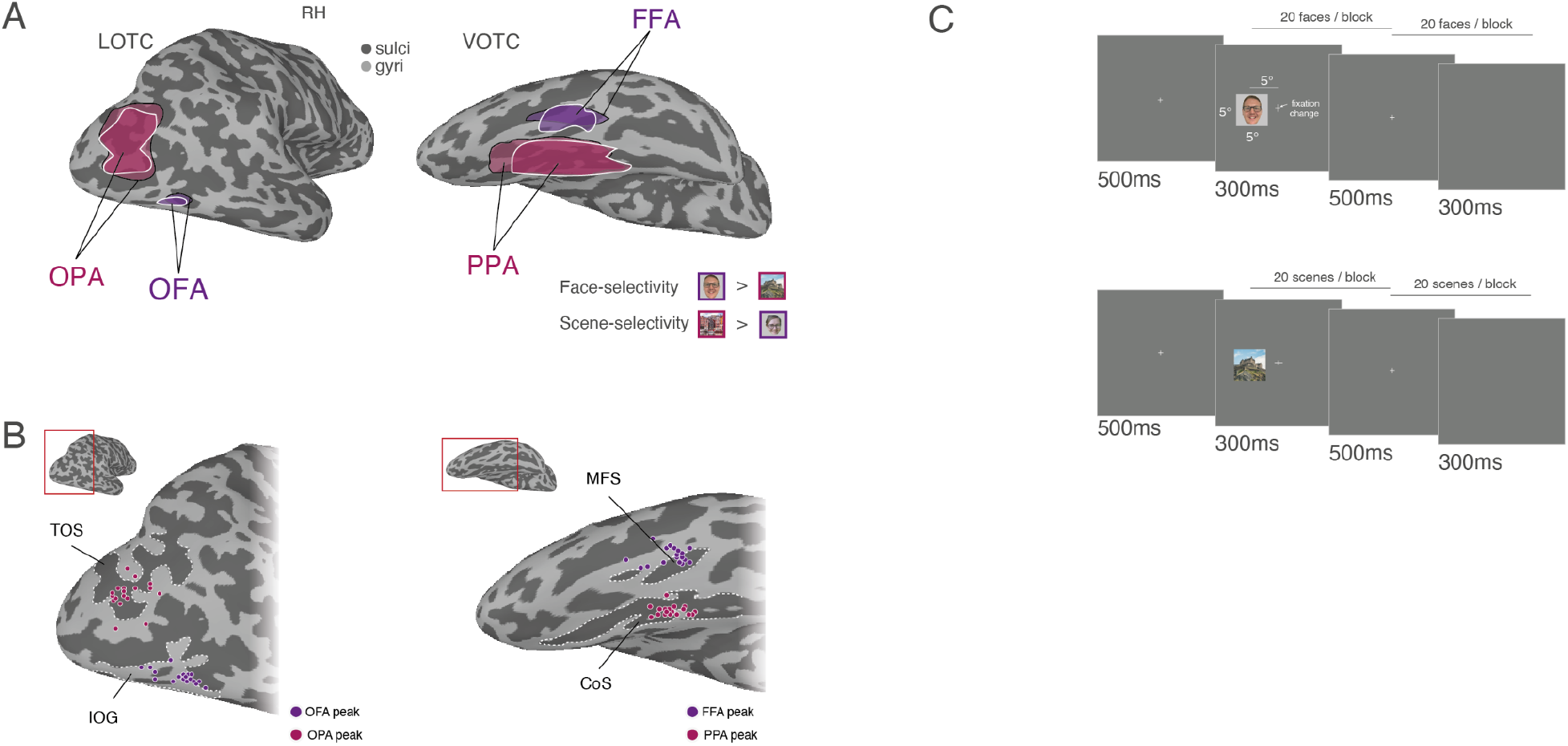
Regions of interest in LOTC and VOTC, individual participant peaks and task schematic. **A,** Lateral (left) and ventral (right) views of a partially inflated right hemisphere are shown (light gray = gyri, dark gray = sulci). Overlaid in pink and outlined in black are the group-based regions of interest for scene-selective Occipital Place Area (OPA) on the lateral surface and Parahippocampal Place Area (PPA) on the ventral surface. Overlaid in pink and outlined in white are the same ROIs (OPA, PPA) but derived from the lateralized experimental runs. Overlaid in purple are the group-based regions of interest for face-selective Occipital Face Area (OFA) on the lateral surface and Fusiform Face Area (FFA) on the ventral surface. Overlaid in purple and outlined in white are the same ROIs (OFA, FFA) but derived from the lateralized experimental runs. All ROIs show a high degree of overlap. Note that ROIs derived from lateralized presentations are not used for analysis, and shown here for illustrative purposes only. **B,** Individual participant ROI peaks. (Left) An enlarged view of the lateral surface of the right hemisphere is shown. Overlaid in pink are the locations of the peak voxel of scene-selectivity in OPA for each individual participant. These show a close correspondence to the transverse occipital sulcus (labelled TOS in B). Overlaid in purple are the locations of the peak voxel of face-selectivity in OFA for each individual participant. These show a close correspondence to the inferior occipital gyrus (labelled IOG in B). (Right) An enlarged view of the ventral surface of the right hemisphere is shown. Overlaid in pink are the locations of the peak voxel of scene-selectivity in PPA for each individual participant. These show a close correspondence to the collateral sulcus (labelled CoS in B). Overlaid in purple are the locations of the peak voxel of face-selectivity in FFA for each individual participant. These show a close correspondence to the mid fusiform sulcus (labelled MFS in B). **C,** Task schematics for face (top) and scene (bottom) blocks. During each 16 s block, 20 images were presented (300ms on / 500ms off) to either the left or right visual fields. During each image presentation one of the fixation cross arms (horizontal or vertical) grew in length. Participants were required to respond via button press which arm was longer. Image exemplars shown here are substitutes and were not shown in the actual experiment.

### 5. fMRI data processing

All anatomical and functional data were pre-processed and analyzed using the Analysis of Functional NeuroImages (AFNI) software (Cox, 1996) (RRID: <u>SCR_005927</u>). Below we outline the preprocessing steps taken for both the initial functional localizer and for the lateralized scene and face sessions.

#### 5.1 Initial functional localizer session

All images were motion-corrected to the first volume of the first run (using the AFNI function *3dVolreg*) after removal of the appropriate dummy volumes to allow stabilization of the magnetic field. Following motion correction, images were detrended (*3dDetrend*) and spatially smoothed (*3dmerge*) with a 5 mm full-width-half-maximum smoothing kernel. Signal amplitudes were then converted into percent signal change (*3dTstat*). To analyze the functional localization data, we employed a general linear model implemented in AFNI (*3dDeconvolve*, *3dREMLfit*). The data at each time point were treated as the sum of all effects thought to be present at that time and the time-series was compared against a Generalized Least Squares (GSLQ) model fit with REML estimation of the temporal auto-correlation structure. Responses were modelled by convolving a standard gamma function with a 16 s square wave for each stimulus block. Estimated motion parameters were included as additional regressors of no-interest and fourth-order polynomials were included to account for slow drifts in the MR signal over time. To derive the response magnitude per category, *t*-tests were performed between the category-specific beta estimates and baseline. The corresponding statistical parametric maps were aligned to the T1 obtained within the same session by calculating an affine transformation (*3dAllineate*) between the motion-corrected EPIs and the anatomical image and applying the resulting transformation matrices to the T1.

#### 5.2 Lateralized face and scene sessions

Functional data from the experimental runs were pre-processed similarly to the pipeline specified above, but differed in the following ways. For each experimental session a mean anatomical image was first computed across the two T1 scans acquired before (Pre) and after (Post) TMS (*3dcalc*). Once pre-processed, all EPI data within a session were then deobliqued (*3dWarp*) and aligned to this mean anatomical image (*align_epi_anat.py*). GLMs were estimated for each run separately (*3dDeconvolve*, *3dREMLfit*) in the unaligned, native volume space, after which the resulting statistical parametric maps were aligned to the mean anatomical image by applying the transformation matrices from the EPI alignment.

#### 5.3 Sampling of data to the cortical surface

In each participant, the pre-processed functional data from all sessions were projected onto surface reconstructions (*3dvol2surf*) of each individual participant’s hemispheres derived from the Freesurfer4 autorecon script (http://surfer.nmr.mgh.harvard.edu/) using the Surface Mapping with AFNI (SUMA) software. The Freesurfer reconstructions were based on the T1s obtained in the localizer session. In order to align the functional data to these surfaces, the mean (Pre-Post) T1 from each TMS/fMRI session was first aligned to the volume used for surface reconstruction (*@SUMA_AlignToExperiment*).

#### 5.4 ROI definitions and analysis

The functional localizer session data was used to define the following ROIs: parahippocampal place area (PPA), occipital place area (OPA), medial place area (MPA, also referred to as RSC), occipital face area (OFA) and fusiform face area (FFA), by overlaying the statistical results of the contrast Scenes versus Faces onto the surface reconstructions of each individual participant, before thresholding (p<0.0001, uncorrected) **(Figure 1A)**. While the broader sampling of categories in our localizer would have allowed for multiple different contrasts to define ROIs, we chose to use this particular one because it matched the planned comparison for our experimental runs (see below). ROIs were defined using the interactive ROI drawing tool in SUMA. ROIs were defined according to both statistical criteria and with respect to accepted anatomical landmarks. For example, PPA was defined as being both scene-selective and located within the collateral sulcus (Weiner et al. 2017, 2018); whereas the FFA was defined as being both face-selective and either overlapping or being lateral to the mid fusiform sulcus (Weiner et al. 2014). No further anatomical or functional constraints were applied. We undertook several steps to ensure our ROI definitions were reliable and consistent with prior work from our own lab (Silson et al. 2015) and others (Weiner et al. 2014, 2017, 2018; Steel et al. 2021). First, we compared the location of these ROIs at the group-level with ones derived from the experimental runs themselves (which employed a lateralized presentation protocol). On average, ROI definitions were highly overlapping **(Figure 1A).** The peak voxels within each ROI (in the right hemisphere) are displayed for all participants in **Figure 1B**. Despite some individual variation, peak voxels adhered to known anatomical landmarks (e.g. all PPA peaks are within the collateral sulcus). Second, we calculated the proportion of our original ROIs that would be included if we had used the contrast of buildings > faces (as opposed to scenes > faces). Although on average ROIs would have been slightly smaller, the vast majority of voxels in our original ROIs would have remained had we chosen this alternative contrast (min proportion across ROIs = 0.70, max proportion across ROIs = 0.95).

Two additional early visual cortex ROIs were defined by projecting a retinotopic atlas (Wang et al. 2015) onto each participant’s surface reconstruction and combining regions V1d, V2d and V3d (for dorsal EVC) and V1v, V2v and V3v (for ventral EVC), respectively. Once defined, the vertices comprising these ROIs were converted to a 1D index of node indices per ROI (*ROI2dataset*), which was subsequently used to extract *t*-statistics for each stimulus category from the three separate TMS/fMRI sessions for each surface node within the ROI (*ConvertDset*). The extracted *t*-statistics were then imported into Matlab (Version R2018B) and averaged across nodes within each ROI.

### 6. fMRI data analysis

#### 6.1 Contralateral and category biases

For each participant and ROI, we computed two types of biases. A Contralateral bias was computed by taking the mean *t*-statistic for the contrast of Contralateral versus Ipsilateral – note, this contrast is collapsed across category (Scenes & Faces). Positive values thus represent a bias for the contralateral visual field, whilst negative values represent an ipsilateral bias. A Face/Scene bias was computed by taking the absolute mean *t*-value for the contrast of Scenes versus Faces – note, this contrast is collapsed across visual field (Contralateral & Ipsilateral). Here, a positive value represents a bias for the preferred category with a negative value representing a non-preferred category bias. These bias measurements for each ROI were taken forward for further analysis.

#### 6.2 ROI statistical analyses

All statistical analyses were performed using the RStudio package (version 1.3.9). Initially, bias values were first averaged across sessions to create a grand-average data set, before subsequent session-specific analyses. In session-specific analyses, bias values were submitted initially to a four-way repeated measures ANOVA with Hemisphere (Left, Right), Surface (Lateral, Ventral), Selectivity (Scene, Face) and Bias (Contralateral, Face/Scene) as within-participant factors. Across all sessions, the main effect of Hemisphere was non-significant thus bias values were collapsed across hemispheres before being submitted to a three-way repeated measures ANOVA with Surface, Selectivity and Bias as within-participant factors (same levels as above). If a significant Surface by Bias interaction was observed, paired *t*-tests were employed to test the strength of the Contralateral versus Face/Scene biases in each ROI separately.

#### 6.3 Whole-brain analysis

To investigate whole brain effects we first calculated whole brain biases (same approach as for the ROIs). Next, we converted these bias indices into estimates of effect size (Cohen’s d: mean Category - mean Contralateral / SD pooled) and projected the result of Contralateral - Category across the cortical surface.

#### 6.4 Within & between bias correlations

First, we split the data into Odd (runs 1&3) and Even (runs 2&4) datasets. Next, in each ROI we pooled biases across hemispheres, resulting in 36 data points per bias. The partial correlation (Spearman’s) between splits (reflecting the within-bias correlation) was computed for each ROI (FFA, OFA, PPA, OPA) and bias (Contralateral, Face/Scene) separately, taking into account the average temporal-signal-to-noise (tSNR). Next, we computed the partial correlation between biases (reflecting the between-bias correlation) for each ROI separately, again taking into account the average tSNR.

## Results

### Strength of category and contralateral biases differs between lateral and ventral ROIs

Initially, we sought to compare directly the strength of contralateral and face/scene category preferences within scene- and face-selective regions across LOTC and VOTC, respectively. Before comparing contralateral and category preferences, we first calculated the mean response to all four conditions (ipsilateral scene, contralateral scene, ipsilateral face, contralateral face) in each ROI (**Figure 2)**. As expected, these data demonstrate the presence of both types of bias (contralateral, category) in each ROI. Indeed, each region showed on average larger responses to stimuli in the contralateral visual field, as well as larger responses to its preferred stimulus (scene/face). These categorical preferences were evident whether stimuli were presented in the contralateral or ipsilateral visual fields (**Figure 2A–D**).

**Figure 2:**
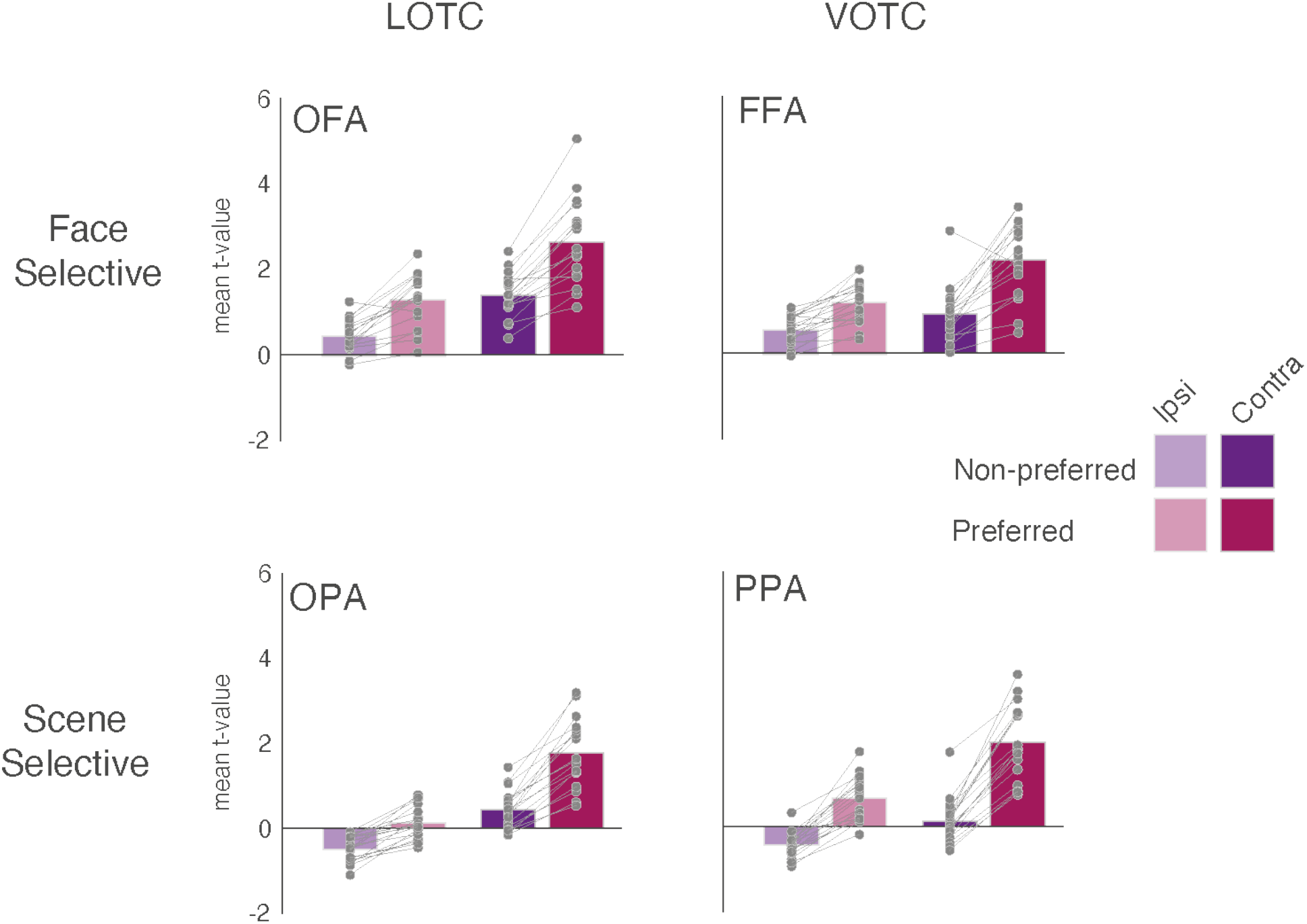
Mean response to all conditions. Bars represent the mean response (t-value versus baseline) for each condition in each ROI (light purple = ipsilateral non-preferred, dark purple = contralateral non-preferred, light pink = ipsilateral preferred, dark pink = contralateral preferred). Face-selective ROIs are plotted on the top row, with scene-selective ROIs on the bottom row. Lateral ROIs are in the left column, ventral ROIs in the right column.

To quantify the strength of the observed category and visual field preferences we computed contralateral and category bias indices for each ROI (see Methods). A series of *t*-tests (against zero = no bias) confirmed that both biases were significantly represented in all ROIs (p<0.001, in all cases).

Having established that all ROIs significantly exhibit both contralateral and category biases simultaneously, we next tested how the relative magnitude of these biases differed between the lateral and ventral pairs of regions. Qualitatively, Figure 2 suggests that responses in LOTC were more strongly influenced by visual hemifield location, while responses in VOTC were more strongly biased towards scene/face category. Specifically, the response to the preferred category in the ipsilateral visual field is stronger than the non-preferred category in the contralateral visual field in VOTC (i.e. the bars follow a ‘sawtooth pattern’), but not LOTC, where the responses is roughly equivalent (in OFA) or always stronger for the contralateral visual field (in OPA). This pattern of results is suggestive of a relatively stronger contralateral bias in LOTC and a relatively stronger category bias in VOTC. To test this, we conducted three-way repeated measures ANOVA with Surface (Lateral, Ventral), ROI (Scene-selective, Face-selective) and Bias (Spatial, Category) as within-participant factors. Below, we first outline the results of these analyses for the average of all three sessions (**Figure 3**), before demonstrating the consistency of these effects in each session, separately **(Figure 4**).

**Figure 3:**
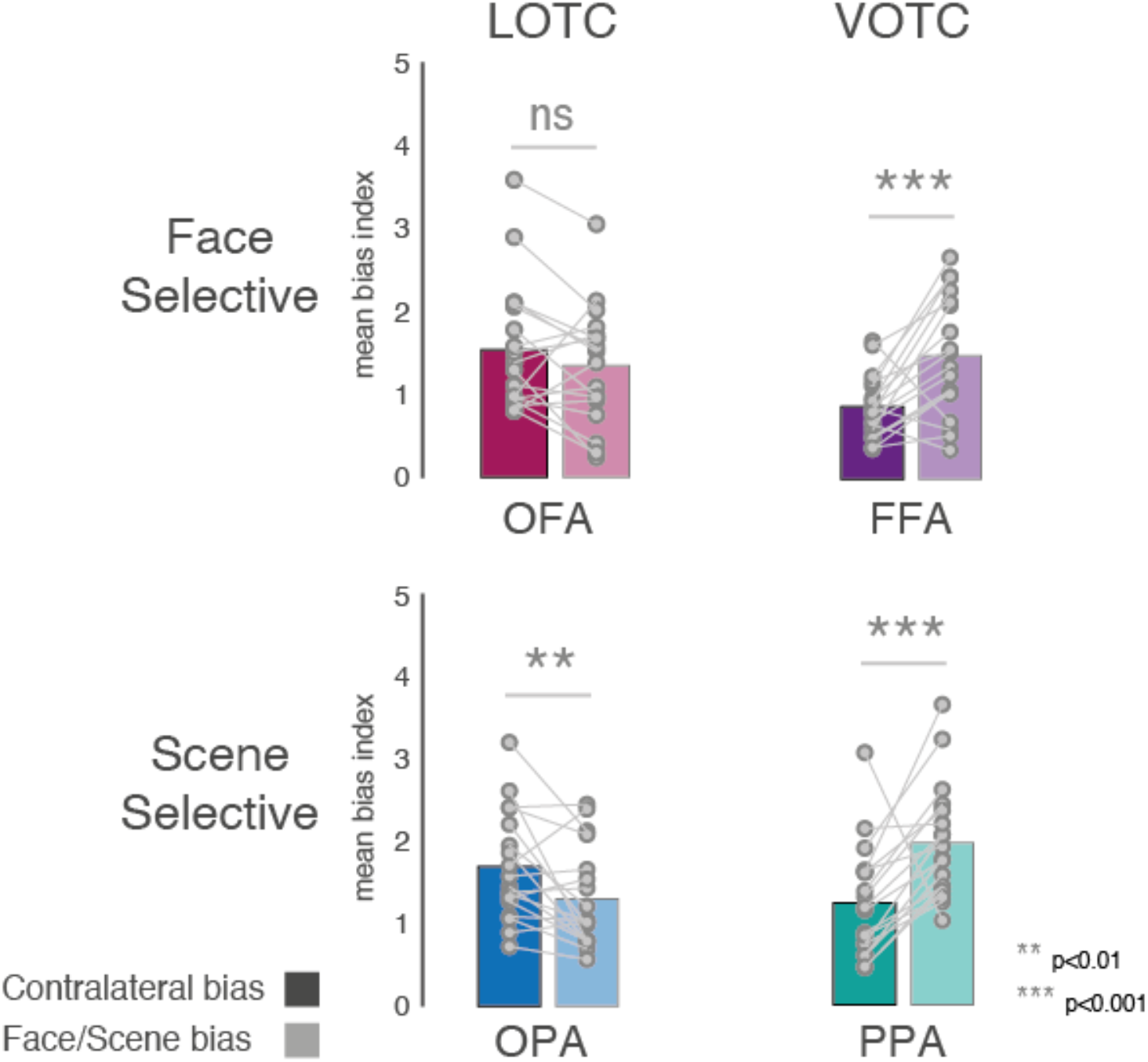
Average contralateral and face/scene category biases. Bars represent the mean contralateral and category biases in each ROI. Individual data points are plotted and linked for each individual and ROI. Contralateral biases are indicated by solid bars, category bias by faded bars. Face-selective ROI = top row, Scene-selective ROIs = bottom row, lateral ROIs = left column, ventral ROIs = right column. On average both lateral ROIs showed a stronger contralateral over category bias (note that this difference was numerically greater in OFA and statistically greater in OPA), whereas both ventral ROIs showed a greater category over spatial bias (ns = non-significant, **p<0.01, ***p<0.001).

**Figure 4:**
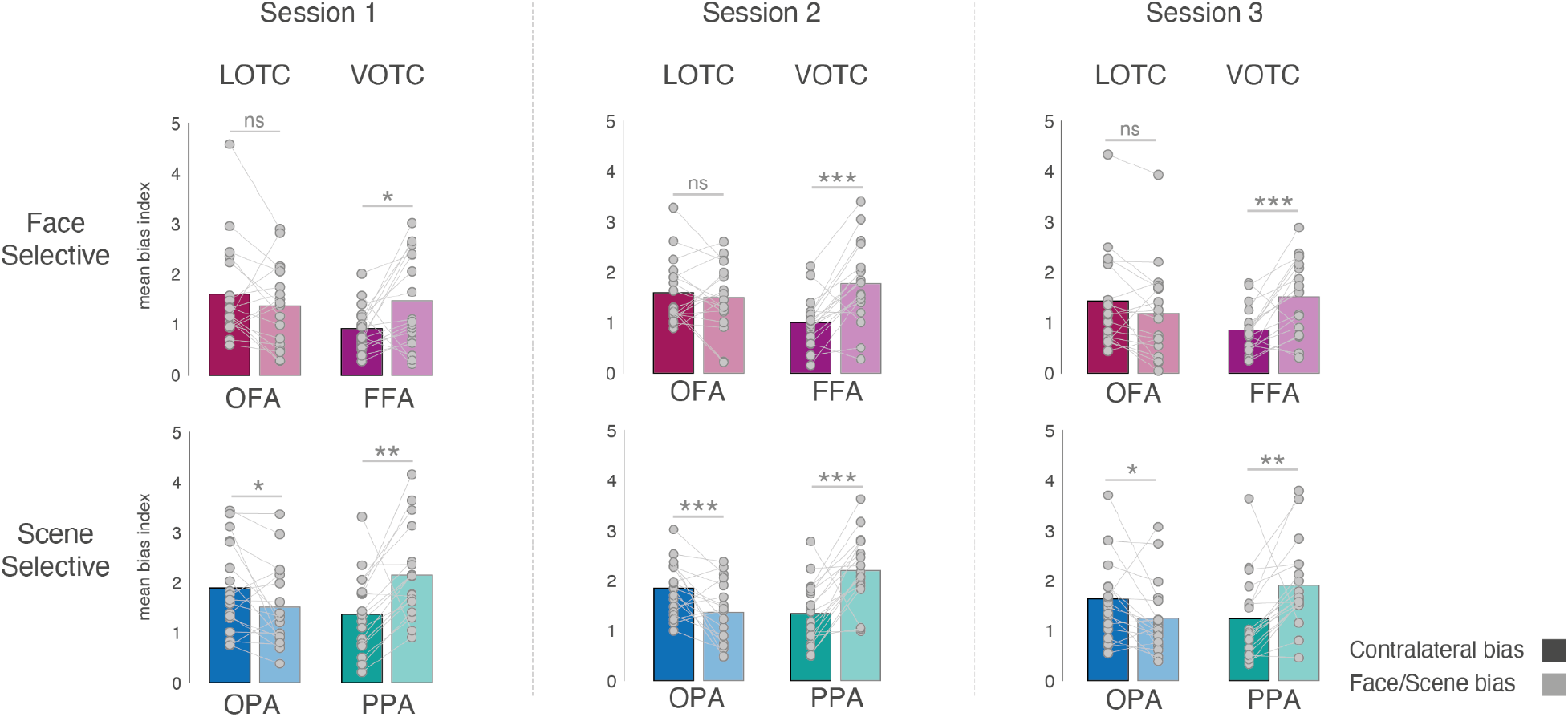
Contralateral and category biases in each individual session. Bars represent the mean spatial and category biases in each ROI and session. Individual data points are plotted and linked for each individual and ROI. In each session, contralateral biases are indicated by solid bars, category biased by faded bars. Face-selective ROI = top row, Scene-selective ROIs = bottom row, lateral ROIs = left column, ventral ROIs = right column. The pattern of biases was extremely similar across sessions. Within each session on average both lateral ROIs showed a stronger contralateral over category bias (note that this difference was numerically greater in OFA and statistically greater in OPA), whereas both ventral ROIs showed a greater category over spatial bias (ns = non-significant, *p=0.05, **p<0.01, ***p<0.001).

Only the main effect of Bias (F(1, 17)=5.09, p=0.04) was significant, reflecting on average larger category over contralateral biases (p>0.05 for all other main effects). The Surface by ROI (F(1, 17)=5.38, p=0.03) interaction was significant, which reflects a larger category bias difference between PPA and OPA compared to FFA and OFA. Crucially, the Surface by Bias interaction was also significant (F(1, 17)=120.31, p=3.92-9; p>0.05, for all other interactions). This interaction reflects a greater contralateral bias in lateral regions, but a greater category bias in ventral regions. To confirm this difference, a series of paired *t*-tests were performed comparing the contralateral versus category bias in each ROI separately (Contralateral vs. Category, OFA: (t(17)=1.46, p=0.16), FFA: (t(17)=3.80, p=0.001), OPA: (t(17)=2.71, p=0.01), PPA: (t(17)=3.74, p=0.001) **(Figure 3)**.

### Category and contralateral biases are consistent across sessions

To examine the consistency of these findings, we next performed the same analyses but for each session separately. These data showed a strikingly consistent pattern across all three sessions, with a significant Surface by Bias interaction present in each case (**Figure 4,** see Supplementary Material for full statistical breakdown).

In addition, we pooled bias values for each participant and session to evaluate the consistency of these biases within participants. This resulted in 16 data points per participant and session (4×ROIs, 2×Biases, 2×Hemispheres). Next, we computed the pairwise across-session correlation (Pearson’s r) in each participant separately, before averaging these correlation coefficients across participants **(Figure 5A)**. A series of *t*-tests (against zero) confirmed on average significant correlations between each pair of sessions (Sessions 1:2 t(17)=13.72, p=1.25-10; Session 1:3: t(17)=15.50, p=1.82-11; Sessions 2:3 t(17)=18.13, p=1,47-12). This demonstrates that on average the contralateral and category biases were consistent within participants across sessions.

**Figure 5:**
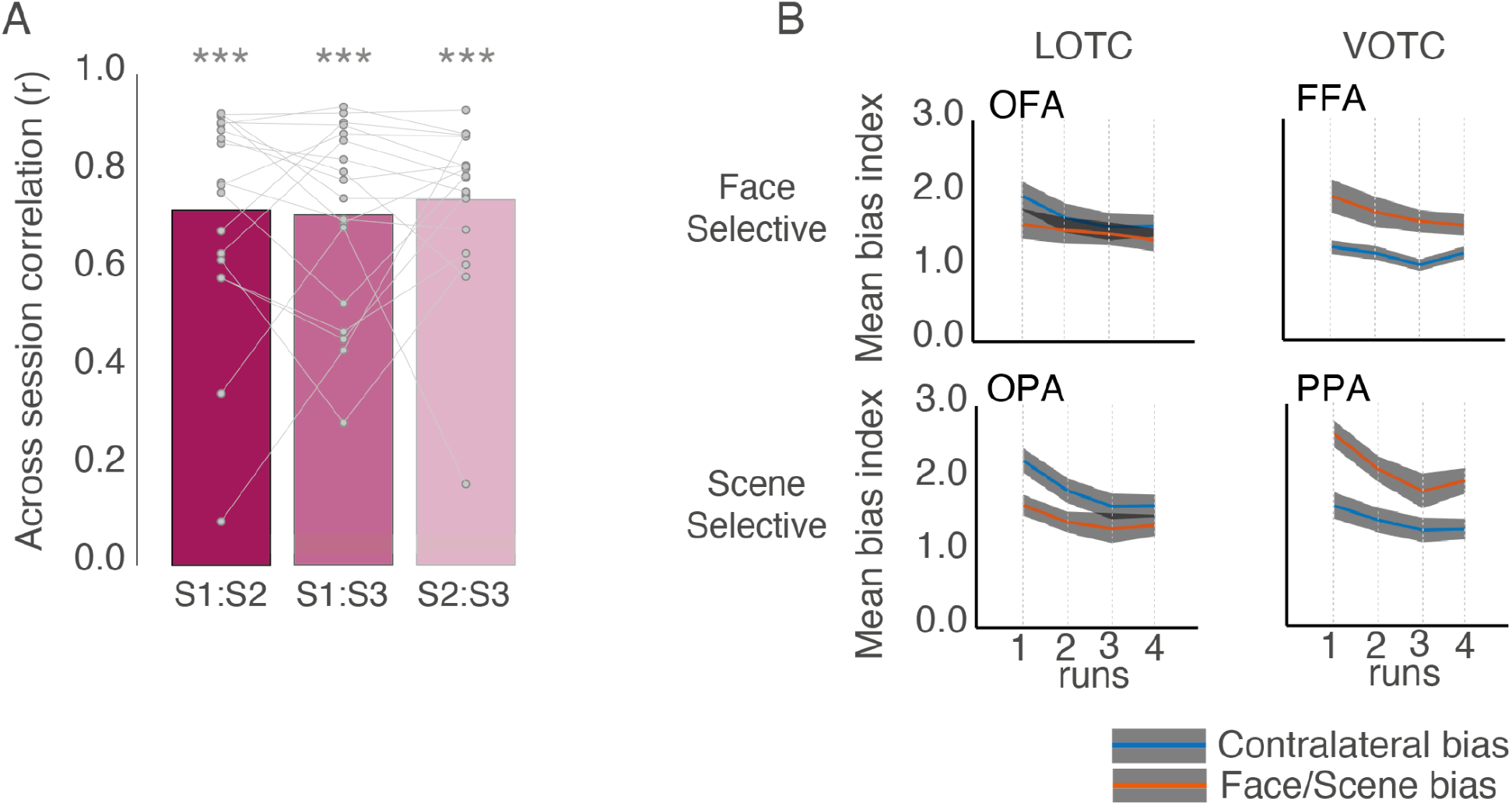
Bias consistency and effect of run. **A**, Bars represent the mean across-session bias correlation (Pearson’s). Individual data points are plotted and linked for each participant. Despite some variability, on average there was a significant correlation between all pairs of sessions. ***p<0.001. **B,** Line plots show the mean (plus s.e.m) bias across runs. Note that due to a non-significant main effect of Session, bias measurements were collapsed across sessions. On average, all ROIs show a general decrease in bias magnitudes across runs, consistent with previous reports reporting overall magnitude decreases across runs. Importantly, this run effect does not alter the relationship between biases within each ROI.

### Category and contralateral biases are consistent across runs

Prior work from our group (Groen et al. 2021) and others (Meshulam and Malach 2016) have highlighted the systematic reduction in fMRI evoked responses that can occur if the same task is performed across multiple repeated fMRI runs. Indeed, our prior work (Groen et al. 2021) demonstrated a widespread effect of run throughout visual cortex during repeated runs of a two-back task involving eight different categories. The analyses thus far were computed on the average biases across runs, but we also looked at the spatial and category biases between runs in each ROI separately **(Figure 5B)**. For each ROI, bias indices were submitted to a three-way repeated measure ANOVA with Session (Session1, Session2, Session3), Run (Run1, Run2, Run3, Run4) and Bias (Spatial, Category) as within-participant factors.

All four ROIs exhibited a significant main effect of Run (*p*<0.05, in all cases) reflecting on average the gradual reduction in response magnitude across successive runs. The main effect of Bias was significant in FFA, OPA and PPA (*p*<0.05), but not OFA (*p*>0.05), which reflects on average a consistently larger contralateral bias in OPA, but a larger category bias in PPA and FFA, respectively. Only in OPA and PPA did we observe a significant Run by Bias interaction (OPA: F(3, 51)=3.25, *p*=0.02; PPA: F(3, 51)=2.87, *p*=0.04), which reflects the tendency for the biases to become more similar across runs in the case of OPA, and a larger difference between the biases in Run 4 as compared to Run 3 in PPA (*p*>0.05, for all other interactions).

These results demonstrate a systematic effect of run on fMRI responses, showing modest interaction with bias strength in some but not all of the ROIs. Importantly, despite the overall reduction in response across runs, the relative magnitude of the biases does not flip in any ROI. That is, the dominance of one bias over the other remains constant in each ROI.

### Stronger within than between biases within participants

Next, we asked to what extent the category and contralateral biases were consistent within participants, and whether or not a stronger category bias might be coupled with a weaker contralateral bias or vice versa. From the traditional/hierarchical viewpoint, category-selectivity and spatial selectivity may trade off against one another, and thus one might predict a negative correlation between biases. Within each ROI we first examined the within-bias similarity before evaluating whether a systematic relationship between biases existed using a split-half analysis (see methods section 6.4). On average within-bias correlations were high (and statistically significant) **(see left panel Figure 6),** reflecting a high level of consistency across independent datasets. Next, we computed the between-bias correlation in each ROI across independent datasets **(see right panel Figure 6)**. Although, on average between-bias correlations were markedly reduced compared to the within-bias measurements, significant correlations remained in OFA (p<0.001), FFA (p=0.03) and OPA (p<0.01), but not PPA (p=0.16). These results show that participants showed both highly reliable contralateral and Face/Scene biases. Moreover, this suggests that within a given participant, these two effects are related: a stronger spatial bias was associated with a stronger category bias, rather than the two effects trading off against one another. It is also worth noting that the between-bias correlations were numerically higher in lateral OFA/OPA than in FFA/PPA - their ventral counterparts.

**Figure 6:**
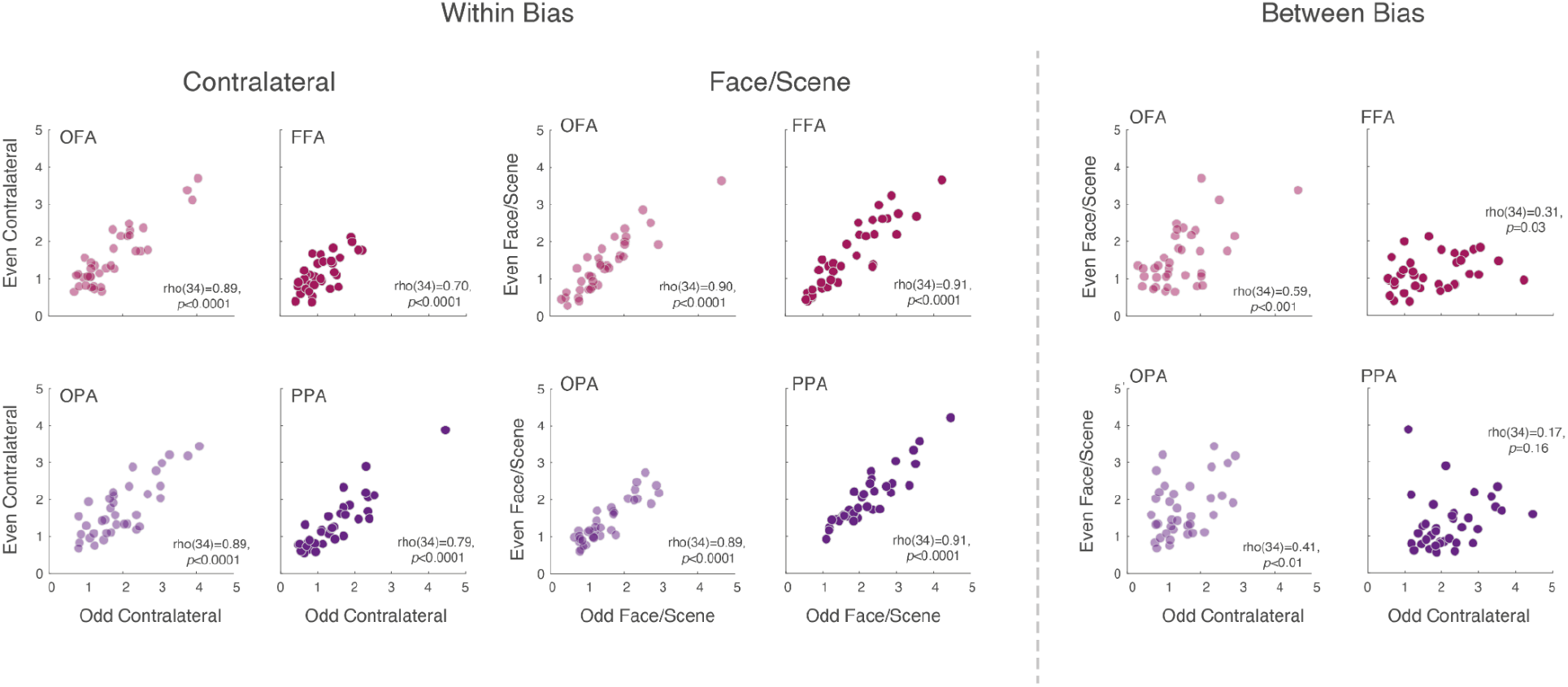
Contralateral and Face/Scene bias relationship: **Left**, Scatter plots show the relationship within the Contralateral and Face/Scene biases across independent datasets within each ROI, while taking into account the tSNR. A significant positive correlation (Spearman’s ρ) was present within all ROIs for both biases. **Right,** Scatter plots show the relationship between the Contralateral and Face/Scenes biases across independent datasets within each ROI. Although weaker than the within-bias correlations, these were significant in OFA, FFA and OPA but not PPA.

### Contralateral bias dominates in early visual cortex

Across all three sessions we observed a consistent Surface by Bias interaction, indicating that a stronger contralateral response laterally but a stronger scene/face category response ventrally. Given that lateral and ventral regions of OTC fall directly anterior of dorsal and ventral early visual regions (V1-V3), respectively, we calculated the mean contralateral and category biases in these regions for comparison. Despite significant contralateral and category biases in these ROIs (min contralateral t=9.45, max contralateral p=3.47-8; min category t=9.92, max category p=1.73-8), the contralateral biases were consistently larger, as expected. A two-way repeated measures ANOVA with ROI (V1-V3d, V1-V3v) and Bias (Spatial, Category) revealed only a significant main effect of Bias (F(1, 17)=28.05, p=5.92-5), which reflects the expected larger contralateral biases in both ROIs (p>0.05, in all other cases). Thus, unlike lateral and ventral scene- and face-selective regions, the magnitude of the contralateral biases are equivalent in dorsal and ventral early visual cortex (**Figure 6B**).

### Category bias dominates in scene-selective MPA

Although our main ROI focus was on the scene- and face-selective regions of the lateral and ventral surfaces, we also calculated the contralateral and category biases in scene-selective Medial Place Area/Retrosplenial complex (MPA/RSC) located on the medial surface of OTC (Figure 5D). Whilst both biases were found to be significantly present (Contralateral: t(17)=8.34, p=2.04-7; Category: t(17)=5.70, p=2.60-5), the category bias was significantly larger than the contralateral bias (Spatial v Category: t(17)=4.64, p=2.29-4). The larger category bias in MPA follows a similar pattern to PPA, although the category advantage is much larger.

### Contralateral and category biases vary gradually across the cortical surface

While the ROI analyses show strong contralateral and category biases, they are limited to the specific choice of categories used to map them (i.e. faces/scenes). In order to explore the relationship between contralateral and category biases outside of our initial ROIs, we computed the whole-brain difference in effect size (Cohen’s d) for each bias and projected the group average maps onto the cortical surface (**Figure 8A**). This qualitative and exploratory analysis reveals three main patterns of results.

**Figure 7:**
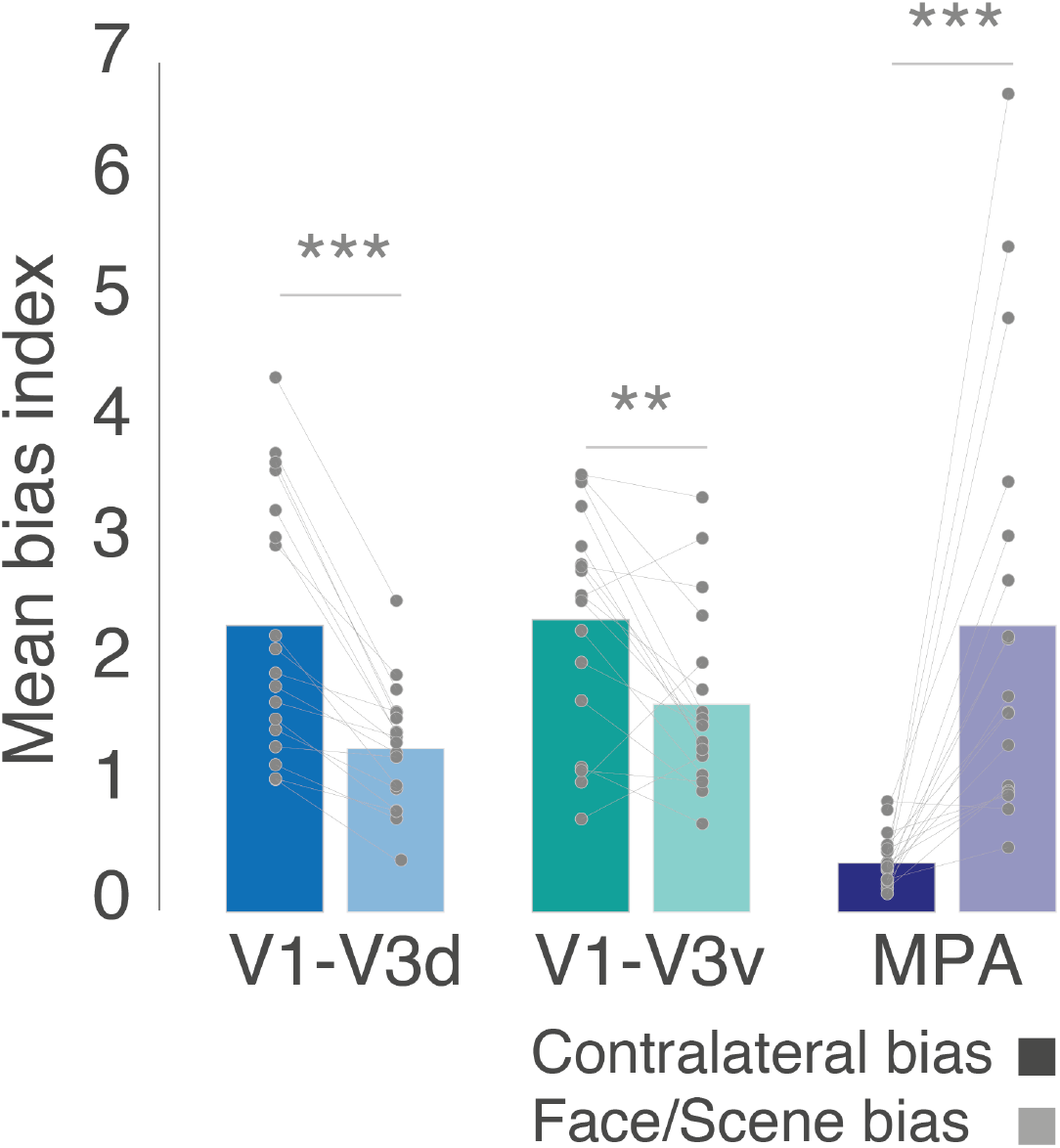
Contralateral and Face/Scene biases in control ROIs: Bars show the mean contralateral and Face/Scene biases in early visual ROIs (V1-V3d, V1-V3v) and a third scene-selective ROI (MPA/RSC). Individual data points are plotted and linked for each individual and ROI. Both V1-V3d and V1-V3v showed an expected greater contralateral bias, whereas MPA showed a greater Face/Scene bias. **p<0.01, ***p<0.001

**Figure 8:**
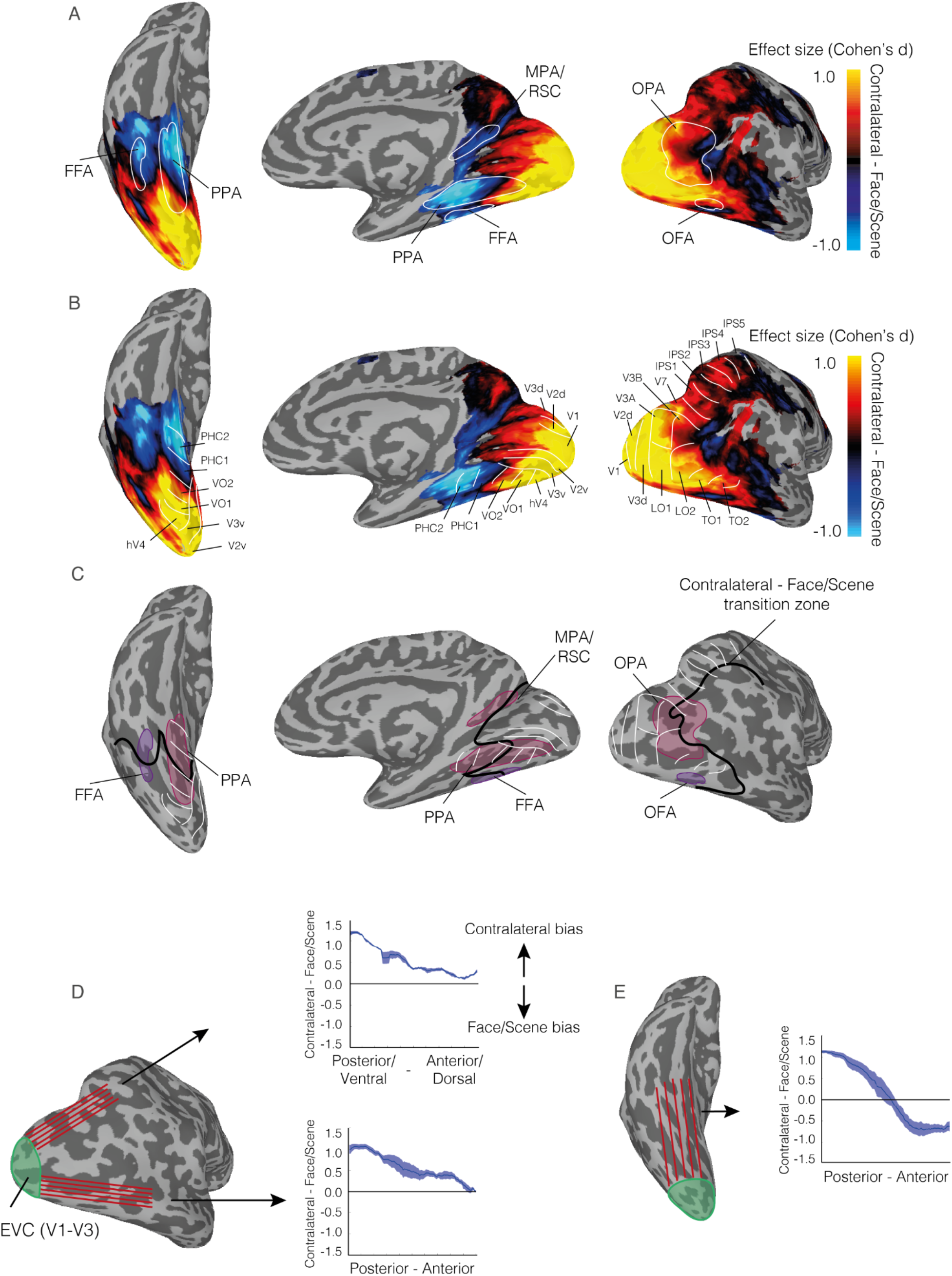
Direct comparison of contralateral and face/scene biases across the cortical surface. **A,** The group average difference in effect size (Cohen’s d) is overlaid onto ventral (left), medial (middle) and lateral (right) view of the right hemisphere. Hot colours represent larger contralateral bias effect sizes with cold colours representing larger face/scene bias effect sizes. The group average ROIs for FFA, PPA, OFA, OPA and MPA/RSC are also overlaid in white. Face/scene bias becomes more predominant in anterior relative to posterior sections of most ROIs. **B,** Same as in A, but with the borders of multiple retinotopic maps (defined using a probabilistic atlas Wang et al., 2014) overlaid. On the lateral surface, these retinotopic borders show a close correspondence to areas showing a greater contralateral bias. On the ventral surface, the border between retinotopic maps VO2 / PHC1 show a close correspondence to the transition zone (black) between the two biases. **C**, White-lines represent the locations of each retinotopic map (max probability from Wang et al., 2014). The group average ROIs are overlaid in pink (OPA/PPA) and purple (OFA/FFA) and the transition zone between contralateral bias and face/scene bias is overlaid in black. On the lateral surface, this transition zone closely follows the anterior border of the retinotopic maps. On the ventral surface, this transition zone cuts across both PPA and FFA and closely matches the border between retinotopic maps VO1 / PHC1. **D**, A lateral view of the right hemisphere is shown. Red-lines represent surface vectors that begin at the anterior border of V3d and project anteriorly into parietal cortex and occipital cortex, respectively. Plots represent the mean bias value (plus sem across subjects) along each vector. Positive values represent larger contralateral bias effect sizes, with negative values representing larger face/scene bias effect sizes. On the lateral surface, although the magnitude of the contralateral biases are reduced anteriorly they remain above the unity line. **E,** A ventral view of the right hemisphere is shown. Red-lines represent surface vectors that begin at the anterior border of V3v and project anteriorly. Unlike on the lateral surface, the magnitude of the spatial bias decreases anteriorly and transitions to represent a stronger face/scene bias more anteriorly.

First, they highlight that the contralateral-to-category biases change smoothly across both the lateral and ventral surfaces, despite differences between surfaces. Second, they highlight how the ‘transition zones’ (where the predominant bias flips) do not map cleanly onto commonly accepted category ROIs on either surface. On the lateral surface, this transition zone nicely aligns with the anterior borders of known retinotopic maps (Wang et al., 2014), but this alignment is less clear ventrally: the borders of PHC1/PHC2 maps clearly overlap categorically biased portions of VTC, whereas the borders of VO1/VO2 clearly overlap contralaterally biased portions of VTC (**Figure 8B**).

Third, notwithstanding the general posterior-anterior gradient present throughout visual cortex, a closer look at how these gradients intersect the retinotopic maps across the lateral and ventral surfaces highlights a distinction between them. Whereas ventrally, there is a clear transition zone running largely medial-lateral in VOTC and corresponding with the border of VO1/PHC1, laterally, the contralateral bias remains largely dominant throughout (**Figure 8C**). Indeed, the contralateral bias persists dorsally all the way into parietal cortex and anteriorly towards the temporal lobe. Interestingly, these data also hint at a potential third gradient that runs from early visual cortex towards the superior temporal sulcus. Here, along this trajectory, there is a more clearly visible transition from contralateral to category that then runs towards the posterior superior temporal sulcus (pSTS).

Together, these whole brain results suggest that the category and contralateral biases observed in our ROI analysis arise from smoothly varying gradients that systematically change from posterior to anterior visual cortex. We find that the contralateral-to-category transition zones in these gradients do not cleanly map onto either a purely retinotopic, atlas-based parcellation nor independently defined category ROIs. Instead, a gradient from contralateral-to-category bias can be observed *within* nearly all of the category-selective ROIs we investigated here. Importantly, the interaction between lateral and ventral surface ROIs that we observed in the ROI analyses seems to reflect qualitative differences in these gradients across these two surfaces, with relatively stronger dominance of contralateral biases throughout the lateral surface compared to the ventral surface.

## Discussion

Here, by using a systematic test of visual field (contralateral - ipsilateral) and category biases (scenes - faces) we demonstrate that while scene- and face-selective regions in LOTC and VOTC exhibit both types of biases, there is a striking difference in the predominant bias from contralateral in LOTC to category in VOTC. These data suggest that category-selective regions in LOTC and VOTC may play different yet complementary roles in visual perception.

### Contralateral and category biases are co-localised

Historically, spatial biases in the form of contralateral representations were considered a hallmark of regions within early visual cortex, for which clear retinotopic maps were established (e.g. V1-V4). In contrast, the identification of category-selective regions more anteriorly throughout LOTC and VOTC, coupled with the lack of evidence for overlapping retinotopic maps at that time, contributed to the idea that these regions exhibited position invariance. Spatial and category biases were thus considered to be largely represented independently. Subsequent fMRI studies, however, revealed such a distinction was overly simplistic (Grill-Spector and Malach 2004). Early fMRI work (Levy et al. 2001; Hasson et al. 2002) demonstrated that face- and scene-selective regions of VOTC overlapped foveal and peripheral visual field representations, respectively. Later, spatial biases in the form of contralateral preferences were identified within several category-selective regions, and more recent work has delineated multiple retinotopic maps that spatially overlap several category-selective regions of LOTC and VOTC. Consistent with prior work (Hemond et al. 2007; MacEvoy and Epstein 2007; Schwarzlose et al. 2008; Chan et al. 2010; Kravitz et al. 2010; Silson et al. 2015; Uyar et al., 2016), our analyses demonstrate that each ROI exhibits both a category bias and simultaneously a contralateral visual field bias. The contemporary view of LOTC and VOTC is thus one whereby category and visual field preferences coexist as opposed to being represented by different regions within the visual hierarchy.

### Contralateral bias dominates laterally, Face/Scene bias dominates ventrally

Our findings extend this prior work by comparing directly the strength of scene/face category and contralateral biases within the ROIs themselves and across visual cortex more broadly. Crucially, we demonstrate that the relative strength of these biases differs between LOTC and VOTC. Specifically, LOTC regions show a stronger contralateral over category bias, whereas ventral regions show a stronger category over contralateral bias. Importantly, the dissociation between LOTC and VOTC was not restricted solely to our face- and scene-selective ROIs. Indeed, although both surfaces showed a general transition from stronger contralateral bias to stronger category bias along the posterior-anterior axis, there remained distinct differences between the two surfaces. On the ventral surface, a clear transition between predominantly contralateral and predominantly category was evident. Interestingly, this clear transition zone showed a close correspondence with the border between retinotopic maps VO2 and PHC1. In contrast, on the lateral surface, the contralateral bias remains relatively dominant throughout, extending dorsally into the parietal cortex and anteriorly in the direction of TO1 and TO2. It is only near the pSTS that the relative strength of these biases becomes equivalent and further flips so that the category bias is stronger. Interestingly, the pSTS is considered a core component of the face-processing network with a preference for dynamic stimuli and has recently been suggested to be part of a third visual processing pathway specialised for social perception (e.g. faces, bodies) (Pitcher and Ungerleider 2021). Furthermore, a recent study investigating differences between lateral and ventral cortex using pRF mapping and diffusion imaging (Finzi et al. 2021) reports differences in spatial sampling between face-selective regions in pSTS versus those in ventral cortex: specifically, lateral STS showed more peripheral spatial biases than ventral regions, and this differences was related to their respective white matter connections with EVC, providing additional evidence for differences between lateral and ventral regions in how they represent visual space. Note that the OFA (referred to as IOG) is grouped alongside more ventral FFA in this study (Finzi et al. 2021).

### Implications for theoretical frameworks of visual processing

The observation of multiple category-selective regions in visual cortex has previously been considered to reflect their relative position within a hierarchical framework (Taylor and Downing 2011). That is, the lateral and more posterior regions were considered the precursor regions to their ventral more anterior counterparts, with for example, face and body parts more strongly represented in lateral OFA and Extrastriate Body Area (EBA), and whole faces and whole bodies more strongly represented in ventral FFA and Fusiform Body Area (FBA). Taken in this context, the finding that lateral regions exhibit a relatively stronger contralateral bias whereas ventral regions exhibit a relatively stronger face/scene bias is consistent with the hierarchical explanation for matched category-selective regions (Taylor and Downing 2011). On the other hand, prior work from our group (Silson et al. 2015) using pRF modelling found no evidence for a significant increase in pRF size in the ventral (i.e. PPA) over lateral (i.e. OPA) scene regions - a hallmark of the visual hierarchy.

An alternative account for equivalently selective regions on the two surfaces (e.g. OPA, PPA) is that they serve different, yet complementary functions. The double dissociation between surface (LOTC, VOTC) and bias (Contralateral, Face/Scene) reported here can be interpreted as consistent with this viewpoint. Prior work by our group (Kravitz et al. 2010; Silson et al. 2015, 2016; Groen et al. 2017) and others (Baldassano et al. 2016; Bonner and Epstein) have discussed potential functional differences between OPA and PPA in terms of biases for the lower and upper visual fields, but here we demonstrate that on average the representation for the contralateral visual field bias in OPA is more dominant than its preference for scenes (versus faces). The spatial overlap between OPA and multiple retinotopic maps reported previously (Nasr et al. 2011; Silson et al. 2016), coupled with the current data for a stronger contralateral bias overall raises the question as to whether defining OPA solely on the basis of a preferential response to scenes is appropriate. A similar question can also be asked of PPA. Our whole-brain analyses highlight that the posterior portion of PPA is predominantly spatially biased (overlapping retinotopic maps VO1 and VO2), whereas the anterior portion of PPA is predominately category biased (overlapping retinotopic maps PHC1 and PHC2). In contrast to the scene-selective regions, neither the OFA nor the FFA show a clear relationship with underlying retinotopic maps despite their contralateral preferences. The strength of these biases also varied within FFA along the posterior-anterior axis with the Face/Scene bias becoming dominant more anteriorly. Indeed, although FFA showed a significant contralateral bias, the whole-brain analyses suggest that the contralateral bias is only dominant at the very posterior border of FFA **(Figure 8A)**. Here, we chose to consider FFA as a single face-selective unit within VOTC and did not separate it further into putative FFA1/FFA2 clusters along the same posterior-anterior axis (Weiner and Grill-Spector 2012; Uyar et al. 2016). Nevertheless, the increasing dominance of the face/scene bias anteriorly we report here is consistent with prior work (Uyar et al. 2016) which showed that FFA2 responded in a less spatially-specific manner than its more posterior counterpart FFA1.

The finding that OPA exhibited an overall stronger contralateral bias is consistent with recent work linking OPA with the coding of navigational affordances (Julian et al. 2016; Bonner and Epstein), such as representing navigational boundaries or available routes of egress within scenes. The lower field bias exhibited by OPA thus makes it ideally placed to undertake such computations. Within OPA itself there appeared a gradient within a stronger spatial bias posteriorly but a stronger category bias anteriorly, and future work will be required to understand the relationship between navigational affordance coding within OPA and the gradient reported here. Another gradient in OPA was reported by (Lescroart and Gallant 2019) who showed evidence for a shift in the representation of openness (open scenes - closed scenes) from posterior to anterior. Again, how the representations of openness interact with the contralateral and category biases reported here requires further investigation.

Finally, the fact that category biases become dominant more anteriorly in VOTC, and to a lesser extent in LOTC, is worth considering within the context of complementary fMRI work that compared perceptual responses with those elicited during episodic memory recall (Silson et al. 2019; Steel et al. 2020; Bainbridge et al. 2021). In general, these studies report a posterior-anterior transition in the locus of activity elicited during perceptual versus mnemonic tasks within scene- and face-selective regions in LOTC and VOTC. In many cases the mnemonically driven responses extended anteriorly beyond the borders of the selectivity-defined ROIs. Whether or not these regions also exhibit spatial and/or category biases is a key goal for future work.

### Consistency of contralateral biases across studies

One limitation of our approach is that we compared only two stimulus categories (faces and scenes), and that is unclear to what extent the bias indices we computed using these two stimulus categories will generalize to other stimulus categories. However, our contralateral bias results are consistent with several prior studies that measured contralateral preferences using other stimulus categories and in other category-selective regions. For example, one study (Hemond et al., 2007) showed object, face and scene stimuli in both the ipsi- and contralateral visual field and measured fMRI responses in object and face-selective regions. As in our data, their study revealed a contralateral bias for all stimulus categories, which was larger in lateral-occipital ROIs (OFA and object-selective LO) compared to ventral ROIs (FFA and posterior fusiform). Interestingly, the spatial bias they report in FFA appears numerically smaller than we report here (their Figure 2A). This might be due to the fact that in Hemond et al., (2007), stimuli were presented at bigger sizes (8×8 degree stimulus windows) and more foveally (~1 degree from fixation), which may have resulted in a relatively reduced spatial bias in foveally-biased FFA. However, another study (MacEvoy & Epstein, 2007) found large contralateral biases (up 50% reduction for ipsi- vs contralateral presentations) for both object and scene stimuli using larger stimuli than ours (9×9 degrees) presented at 1.5 degrees from fixation, while (Chan et al., 2010) found a relatively modest contralateral preference in body-selective regions for stimuli presented 3 degrees from fixation. Importantly, none of these prior studies directly compared the relative strengths of both biases allowing for the identification of the ‘transition zone’ in Figure 7. Since the overwhelming majority of studies on category perception and the underlying representations in the human brain use foveal presentation paradigms, it is currently unclear how much the presence and magnitude of contralateral biases and the transition zone depends on the stimulus category, stimulus size and the exact position of the stimulus within the visual hemifield, and future work is needed to address to what extent the results we report here generalize across stimulus categories and visual field positions (see Uyar et al. 2016 for a relatively recent approach).

### Future directions

One way to investigate the precise relationship between visual field position and stimulus selectivity more systematically in future fMRI studies is to employ a population receptive field (pRF) mapping approach - as done in previous works for several specific category-selective regions (e.g, Kay et al. 2015; Silson et al. 2015) and the ventral and lateral-occipital surfaces more broadly. However, one drawback of this approach is that pRF measurements are typically expressed in terms of fitted model parameters, rather than (differences) in response magnitudes as done here, making it more difficult to compare spatial and category tuning directly against one another. Nevertheless, this approach would make it possible to quantify potentially separate contributions of different pRF properties, such as pRF size and position, to category-selective responses in higher visual cortex more broadly.

Other directions for future research on the relative strength of spatial and category tuning in visual cortex should focus on the role of attention, as well as other visual field biases. In our paradigm, we instructed subjects to fixate and attend to the fixation cross, in order to prevent eye movements to the lateralized stimuli. However, evidence suggests that attention can change spatial tuning in visual cortex, including pRF sizes and position in category-selective regions (Kay et al. 2015). It is unclear whether these changes in spatial tuning affect or interact with the category preference in higher visual cortex regions. Moreover, as mentioned above, category-selective regions in LOTC and VOTC exhibit systematic visual field biases not only along the horizontal meridian (contralaterality) but also the vertical meridian (upper vs. lower visual field), which are likely inherited from how early visual field maps feed into the ventral and dorsal streams (Kravitz et al. 2010, 2013; Silson et al. 2015; Uyar et al. 2016). Prior work on scene-selective regions suggests that these upper and lower biases may serve functional goals, such as navigation (lower field OPA; (Bonner and Epstein)) or facilitate recognition of global scene properties (upper field PPA; (Silson et al. 2015; Uyar et al. 2016)). Whether or not such field biases affect category tuning per se, across different category-selective regions, is currently unclear and needs to be investigated in future studies.

### Conclusion

By directly comparing the strength of contralateral and categorical preference in fMRI responses to laterally presented face and scene stimuli, we demonstrate a dissociation between scene- and face-selective regions within LOTC and VOTC, with a stronger contralateral bias in LOTC but a stronger category bias in VOTC. These patterns were consistent both within individuals and across multiple scanning sessions. Moreover, we highlight that this change in predominant bias was not restricted to our specific ROIs, but extended throughout LOTC and VOTC, respectively. Taken together, these data suggest different, yet complementary roles for equivalently category-selective regions within LOTC and VOTC.

## Declarations

### Availability of data and material

Data will be made available on the OSF: https://osf.io/afkj6/

### Code availability

Code will be made available on the OSF: https://osf.io/afkj6/

### Authors’ contributions

EHS, IIAG and CIB designed the experiment. EHS and IIAG performed the experiment and analyzed the data. EHS, IIAG and CIB wrote the paper.

### Ethics approval

The National Institutes of Health Institutional Review Board approved the consent and protocol. This work was supported by the Intramural Research program of the National Institutes of Health – National Institute of Mental Health Clinical Study Protocols 93-M-0170 (NCT00001360) and 12-M-0128 (NCT01617408).

### Consent to participate

Informed consent was obtained from all individual participants included in the study.

### Consent for publication

Not applicable

## Supplementary Material

### Consistency of contralateral and category biases

#### Session 1

Only the main effect of Category (F(1, 17)=5.13, p=0.03) was significant, reflecting on average larger bias values in scene- over face-selective ROIs (p>0.05 for all other main effects). The Surface by Category (F(1, 17)=5.19, p=0.03) interaction was significant, again reflecting a larger difference in category bias between PPA and OPA. The Surface by Bias interaction (F(1, 17)=93.93, p=2.44-8) was also significant and reflects on average a greater contralateral bias laterally but a greater category bias ventrally (p>0.05, for all other interactions). A series of paired *t*-tests were performed comparing the contralateral versus category bias in each ROI separately (OFA: Contralateral v Category (t(17)=1.28, p=0.21), FFA: Contralateral v Category (t(17)=2.50, p=0.02), OPA: Contralateral v Category (t(17)=2.10, p=0.04), PPA: Contralateral v Category (t(17)=3.53, p=0.002) **(Figure 4A)**.

#### Session 2

Only the main effect of Bias (F(1, 17)=6.85, p=0.01) was significant, reflecting on average larger Category over Contralateral biases across ROIs (p>0.05, for all other main effects). The Surface by Category (F(1, 17)=4.48, p=0.04) interaction was significant, again reflecting a larger difference in category bias between PPA and OPA, but crucially so was the Surface by Bias interaction (F(1, 17)=137.11, p=2.10-9), (p>0.05, for all other interactions). Again, the Surface by Bias interaction is driven by a greater contralateral bias laterally, but a greater category bias ventrally (OFA: Contralateral v Category (t(17)=0.69, p=0.49), FFA: Contralateral v Category (t(17)=4.59, p=0.0002), OPA: Contralateral v Category (t(17)=2.197, p=0.008), PPA: Contralateral v Category (t(17)=4.28, p=0.0005) **(Figure 4B)**.

#### Session 3

Only the main effect of Category (F(1, 17)=4.76, p=0.04) was significant, reflecting on average larger bias values in scene- over face-selective ROIs (p>0.05) for all other main effects). Only the Surface by Bias (F(1, 17)=63.65, p=3.78-7) interaction was significant (p>0.05, for all other interactions). Again, the Surface by Bias interaction is driven by a greater contralateral bias laterally, but a greater category bias ventrally (OFA: Spatial v Category (t(17)=1.90, p=0.07), FFA: Spatial v Category (t(17)=3.90, p=0.001), OPA: Spatial v Category (t(17)=2.12, p=0.04), PPA: Spatial v Category (t(17)=2.88, p=0.01) **(Figure 4C)**.

